# Shrinking in the dark: Parallel endosymbiont genome erosions are associated with repeated host transitions to an underground life

**DOI:** 10.1101/2023.09.11.557272

**Authors:** Perry G. Beasley-Hall, Yukihiro Kinjo, Harley A. Rose, James Walker, Charles S. P. Foster, Toby G. L. Kovacs, Thomas Bourguignon, Simon Y. W. Ho, Nathan Lo

**Affiliations:** School of Life and Environmental Sciences, University of Sydney, Sydney, New South Wales 2006, Australia; School of Biological Sciences, University of Adelaide, Adelaide, South Australia 5000, Australia; Okinawa Institute of Science & Technology Graduate University, 1919-1 Tancha, Onna-son, Okinawa 904-0495, Japan; Australian Government Department of Agriculture Water and Environment, Canberra 2601, Australia; University of New South Wales, Sydney, New South Wales 2052, Australia

**Keywords:** parallel evolution, *Blattabacterium*, gene loss, endosymbiosis, molecular evolution, subterranean habitats

## Abstract

Microbial symbioses have had profound impacts on the evolution of animals. Conversely, changes in host biology may impact the evolutionary trajectory of symbionts themselves. *Blattabacterium cuenoti* is present in almost all cockroach species and enables hosts to subsist on a nutrient-poor diet. To investigate if host biology has impacted *Blattabacterium* at the genomic level, we sequenced and analysed 25 genomes from Australian soil-burrowing cockroaches (Blaberidae: Panesthiinae) which have undergone at least seven independent subterranean transitions from above-ground, wood-feeding ancestors. We find at least three independent instances of genome erosion have occurred in *Blattabacterium* strains exclusive to Australian soil-burrowing cockroaches. Such shrinkages have involved the repeated inactivation of genes involved in amino acid biosynthesis and nitrogen recycling, the core role of Blattabacterium in the host-symbiont relationship. The most drastic of these erosions have occurred in hosts thought to have transitioned underground the earliest relative to other lineages. As *Blattabacterium* is unable to fulfil its core function in such host groups, our findings suggest soil-burrowing cockroaches must acquire these nutrients from novel sources. Our study represents one of the first cases, to our knowledge, of parallel host adaptations leading to concomitant parallelism in their mutualistic symbionts, further underscoring the intimate relationship between these two partners.

## 1 BACKGROUND

Microbial symbioses play a critical role in the biology of animals. Symbiotic bacteria perform a wide variety of crucial functions for their hosts, ranging from contributions to metabolism and digestion to prey location using bioluminescence (Degnan et al., 2005; Lundgren and Lehman, 2010; Rio et al., 2016; Ruby, 1996). The close relationship between hosts and their symbionts means that adaptations by the former to changing environments may drive evolutionary change in the latter. The degree to which a symbiont evolves when faced with a shift in host biology depends on the level of intimacy the two partners share.

The phenomenon of parallel evolution, the independent evolution of traits stemming from similar initial conditions (Stern, 2013), provides a useful natural experiment to examine the effects of host adaptation on their symbionts. The question as to whether parallel evolution in hosts is reflected in repeated, predictable changes in their symbionts has been addressed in a limited number of host taxa with a focus on gut symbionts. Parallel genomic changes in the growth-promoting gut symbiont of *Drosophila* have been linked to the nutrition of the host (Martino et al., 2018). In sticklebacks, parallel shifts in allele frequencies linked to specific diets might have led to parallelism in the gut microbiome (Rennison et al., 2019). However, instances of probable host-symbiont parallelism can also often defy expectations due to factors like host environment and genetic variation and, in some cases, because the pair have experienced divergent selective pressures. Such patterns have been observed in guppies and lake whitefish, with gut microbiota communities varying between species or microhabitats, but not in a parallel manner (Sevellec et al., 2018; Sullam et al., 2015).

Obligate intracellular symbionts (hereafter endosymbionts) have highly intimate relationships with their hosts, typically being inherited vertically through the eggs and remaining with their hosts throughout their entire lives. It is among these symbionts that we might expect parallel shifts in host biology to lead to parallel changes in symbiont biology. While instances of convergent or parallel evolution have been demonstrated in endosymbionts (Degnan et al., 2005; Manzano-Marín et al., 2016; Sloan et al., 2014), there has been a relative lack of studies on the effects of host parallel evolution on such microbes.

Aside from those in the Hemiptera, cockroaches are some of the best-studied invertebrates in the context of endosymbiosis. Unique to the group is *Blattabacterium cuenoti* (hereafter *Blattabacterium*), found in almost all species apart from the cave-adapted family Nocticolidae and all termites except *Mastotermes darwiniensis* (Evangelista et al., 2019; Kinjo et al., 2018; Lo et al., 2007). *Blattabacterium* occurs in the fat body tissue and is transmitted transovarially, playing a role in provisioning nutrients to hosts from nitrogenous wastes, such as urea and ammonia (Brooks, 1970; Brooks and Richards, 1955; Sabree et al., 2009). As it is strictly vertically transmitted, *Blattabacterium’s* phylogeny accurately recapitulates the ∼200 million years (My) of its host’s evolutionary history (Beasley-Hall et al., 2021; Evangelista et al., 2019; Kinjo et al., 2021, 2018).

We have recently discovered a striking instance of parallelism in soil-burrowing cockroaches endemic to Australia regarding the repeated acquisition of specialised burrowing behaviour from log-dwelling ancestors (Blaberidae: Panesthiinae). Soil-burrowing cockroaches— previously considered their own subfamily, “Geoscapheinae”—construct permanent underground burrows in which they care for their young and are large, long-lived insects with lifespans of up to almost a decade (Rugg and Rose, 1991). These lineages appear to have acquired such traits at least seven times in parallel from above-ground, wood-feeding ancestors driven by aridification of the Australian continent that began in the early Miocene (Beasley-Hall et al., 2021; Lo et al., 2016; Maekawa et al., 2003; Martin, 2006). These species provide an ideal test case for whether parallel changes in host biology have led to parallel changes in their endosymbionts.

To assess this, here we sequenced 25 additional *Blattabacterium* genomes from soil-burrowing cockroaches and their Australian wood-feeding ancestors to assess two competing hypotheses: 1) that parallel evolution in hosts might not necessarily lead to parallel changes to the endosymbiont genome, as has been found in the gut symbionts of host taxa of other species, vs. 2) parallel evolution in hosts will impart similar pressures on endosymbionts in each case, leading to repeated changes to the endosymbiont genome that are themselves analogous. To do this, we assessed gene loss, pseudogenisation, and shifts in selective pressures.

## 2 METHODS

### 2.1 Host taxon sampling and *Blattabacterium* genome assemblies

Twenty-five representatives of the soil-burrowing cockroaches and Australian lineages of the wood-feeding Panesthiinae (Table S1) were selected to maximise sampling from major clades that represent instances of parallel evolution (Beasley-Hall et al., 2021; Lo et al., 2016). Outgroup sequence data were obtained from GenBank, representing strains from other subfamilies within the Blaberidae (Table S2). SPAdes v.3.12.0 (Bankevich et al., 2012), TCSF-IMRA v.2.7.1 (Kinjo et al., 2015), Chromosomer v.0.1.3 (Tamazian et al., 2016), and Pilon v.1.22 (Walker et al., 2014) were used to de *novo* assemble Blattabacterium genomes following shotgun sequencing of host fat body tissue. MetaPhlAn v2.0 (Truong et al., 2015) was used to detect the potential presence of secondary symbionts. Complete information about sequencing and assembly is available in the electronic supplementary material.

### 2.2 Annotation, pseudogene detection, and selection analyses

Bacterial genome annotation was performed using Prokka v.1.14.5 (Seemann, 2014) and a custom BLAST protein database constructed from all *Blattabacterium* genomes deposited via GenBank at the time of writing (Table S2). Gene loss was assessed using Prokka annotations and by eye. Orthologous genes (OGs) were inferred using OMA v2.5.0 (Altenhoff et al., 2021) and pseudogene detection was performed on OGs using Pseudofinder (Syberg-Olsen et al., 2022). Tests for signatures of relaxed and positive selection were conducted using the codeml program in PAML v4 (Yang, 2007). *Blattabacterium* strains originating from soil-burrowing hosts were selected as lineages of interest (foreground branches) for the analysis. We used branch models (one ratio [M0] *vs*. two ratio [M1] model) and branch-site models (model A1 vs. A) to assess positive vs. relaxed selection, respectively, following (Mitterboeck et al., 2017). A more detailed description of our selection analyses is available in the electronic supplementary material.

## 3 RESULTS

We sequenced 25 genome sequences of *Blattabacterium* strains associated with Australian soil-burrowing and wood-feeding lineages of the Panesthiinae (Table S1). Genome sizes ranged from 611.1 to 634.8 kb and 629.3 to 635.5 in soil-burrowing and wood-feeding hosts, respectively (Table 1). The GC content of our genomes was between 25.3% and 26.3%. We observed complete synteny between our genomes and all published *Blattabacterium* strains from the Panesthiinae and, as with other *Blattabacterium* strains from the subfamily (Kinjo et al., 2018), the plasmid encoding the ribonucleotide reductase β subunit has been integrated into the chromosome.

**TABLE 1.** Genome features of *Blattabacterium* strains sequenced here. Distinct populations of host taxa are indicated where applicable. Counts of genome features refer to Prokka annotations.

| Host species | Strain | Size (kb) | GC% | CDS | rRNAs | tRNAs | tmRNAs | Scaffolds |
| --- | --- | --- | --- | --- | --- | --- | --- | --- |
| *Geoscapheus crenulatus crenulatus* | BGCC | 630.9 | 25.9 | 587 | 3 | 34 | 1 | 7 |
| *G. dilatatus* (Charleville) | BGDch | 613.8 | 25.3 | 568 | 3 | 33 | 1 | 2 |
| *G. dilatatus* (Gilgandra) | BGDgi | 613.9 | 25.3 | 570 | 3 | 33 | 1 | 1 |
| *G. dilatatus* (Mitchell) | BGDmi | 623.2 | 25.7 | 564 | 3 | 33 | 1 | 5 |
| *G. dilatatus* (Patchewollock) | BGDpa | 616.5 | 25.3 | 571 | 3 | 33 | 1 | 11 |
| *G. dilatatus* (Renmark) | BGDre | 614 | 25.3 | 570 | 3 | 33 | 1 | 4 |
| *G. dilatatus* (Walpeup) | BGDwa | 614 | 25.3 | 568 | 3 | 33 | 1 | 3 |
| *G. dilatatus* (Yenda) | BGDye | 613.9 | 25.3 | 569 | 3 | 33 | 1 | 4 |
| *G. robustus* | BGR | 611.1 | 25.4 | 564 | 3 | 31 | 1 | 1 |
| *G. woodwardi* | BGW | 621.9 | 25.9 | 564 | 3 | 33 | 1 | 16 |
| *Macropanesthia kinkuna* | BMK | 632.2 | 26.1 | 597 | 3 | 34 | 1 | 117 |
| *M. lithgowae* | BML | 631.6 | 25.8 | 586 | 3 | 34 | 1 | 15 |
| *M. mutica* | BMM | 625.5 | 26.2 | 586 | 3 | 33 | 1 | 1 |
| *M. rothi* | BMRO | 634.8 | 26.3 | 581 | 3 | 34 | 1 | 43 |
| *Neogeoscapheus dahmsi* | BND | 623.8 | 25.8 | 574 | 3 | 33 | 1 | 12 |
| *N. hanni* | BNH | 624.7 | 26.1 | 583 | 3 | 34 | 1 | 43 |
| *Panesthia australis* | BPAU | 632.4 | 25.9 | 578 | 3 | 34 | 1 | 11 |
| *P. obtusa* | BPO | 634.6 | 26.2 | 578 | 3 | 34 | 1 | 10 |
| *P. sloanei* (Mossman Gorge) | BPSmg | 633.2 | 26.3 | 583 | 3 | 34 | 1 | 6 |
| *P. sloanei* (Mt Lewis) | BPSml | 634 | 26.4 | 591 | 3 | 34 | 1 | 24 |
| *P. tryoni tegminifera* | BPTT | 635.5 | 26.1 | 590 | 3 | 34 | 1 | 24 |
| *P. tryoni tryoni* (Kroombit) | BPTTk | 632.3 | 25.9 | 584 | 3 | 34 | 1 | 7 |
| *P. tryoni tryoni* (Eungella) | BPTTeu | 635.4 | 26.1 | 586 | 3 | 33 | 1 | 6 |
| *P. tryoni tryoni* (Sarina) | BPTT | 631.5 | 26 | 581 | 3 | 33 | 1 | 27 |
| *Parapanesthia gigantea* | BPG | 629.3 | 26.1 | 580 | 3 | 33 | 1 | 33 |

Endosymbionts of soil-burrowing hosts examined here have experienced consistent, extensive gene losses compared to str. BPAY (Figure 1). In contrast, we did not observe such losses in wood-feeding host species. Repeated losses of genes required for amino acid (AA) biosynthesis have occurred almost exclusively in soil-burrowing host-derived endosymbionts. Each of these strains has lost, or experienced pseudogenization in, at least one gene involved in the biosynthesis pathway of leucine, isoleucine, valine, phenylalanine, tyrosine, and tryptophan. We observed little variation in gene inventories in endosymbiont strains sequenced from conspecific hosts. The most extensive losses correspond to strains from Geoscapheus dilatatus and Geoscapheus robustus. These lineages have lost the entire pathway necessary for the biosynthesis of branched-chain amino acids (isoleucine, leucine, valine) and almost all the genes required for tryptophan biosynthesis except for *trp*E.

**FIGURE 1.**
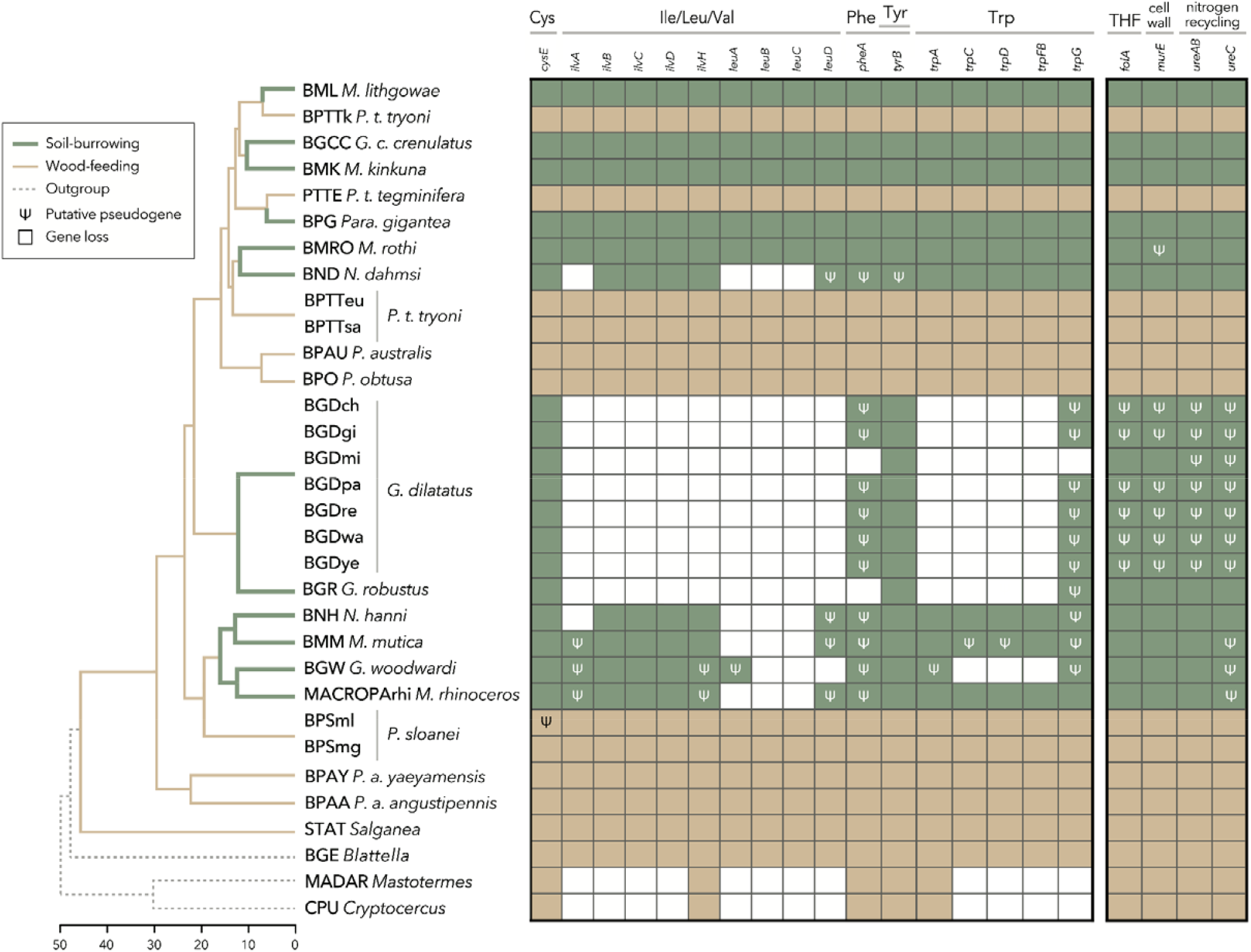
The loss and degradation of genes in *Blattabacterium* involved in the biosynthesis of amino acids (left box) or vitamins and metabolites (right) occurs almost exclusively in soil-burrowing, but not wood-feeding, host lineages. Only genes with evidence of extensive loss or pseudogenisation compared to str. BPAY are shown here. Ingroup relationships were sourced from both host and symbiont genomic data (Beasley-Hall et al., 2021). All amino acids in the left box are essential AAs apart from tyrosine. Host divergence times are shown at the axis on the bottom left in millions of years. Outgroup relationships (dotted lines) follow (Djernæs et al., 2020) and are not time calibrated. THF refers to tetrahydrofolic acid and “cell wall” refers to peptidoglycan biosynthesis.

We detected events of putative pseudogenization in 201 of 715 orthologous groups (Supplementary File S1) prior to manual curation, based either on the predicted fragmentation of a single gene into multiple OGs or a gene being truncated compared to closely related strains. Notably, over 70% of these putative pseudogenization events occurred in *Blattabacterium* strains derived from soil-burrowing hosts (Supplementary File S1). Pseudogenized genes included those involved in the biosynthesis of amino acids, vitamins, cell wall biosynthesis, and nitrogen assimilation (Supplementary File S1; Figure 1). We did not observe any consistent gene losses or pseudogenization events in wood-feeding host-derived strains related to biosynthesis genes.

One of the most striking instances of pseudogenisation in our dataset involves the alpha and gamma subunits of urease (*ureC, ureAB*), involved in the conversion of urea to ammonia, a crucial step in the nitrogen recycling pathway of Blattabacterium. These two genes have been seemingly rendered non-functional on at least two separate occasions, once in strains from *Geoscapheus dilatatus* and *robustus*, and again in str. BMM (*Macropanesthia mutica*) and BGW (*Geoscapheus woodwardi*) (Supplementary File S2; Figure 1).

No large, non-*Blattabacterium* contigs were consistently detected in any sample using MetaPhlAn2, suggesting the absence of secondary symbionts in the fat body tissue of hosts and in agreement with previous work (Kinjo et al., 2018). However, this does not rule out the presence of secondary symbionts outside of the fat body, and we note that similar microscopy work has not been conducted on Australian members of the Panesthiinae. We found no evidence for relaxed selection amongst *Blattabacterium* lineages after correcting for multiple tests (Supplementary File S3). Twelve genes showed evidence of positive selection using branch-site models in PAML including *aceF, argE/dapE, dnaG, glnS, guaB, murB, purC, sucB, and tadA* (Supplementary File S4; *p*-values ranging from 0.001029 to 2.35E-09), involved in the biosynthesis of products including amino acids, metabolites, and peptidoglycan.

## 4 DISCUSSION

### 4.1 Parallel and repeated genome erosions in *Blattabacterium* strains are exclusive to soil-burrowing cockroaches

Our results demonstrate that parallel gene losses have almost exclusively occurred in *Blattabacterium* strains found in soil-burrowing, but not wood-feeding, hosts of the subfamily Panesthiinae. Extensive losses are all but restricted to genes involved in the biosynthesis of amino acids (hereafter “gene losses”), whereas pseudogenization is more widespread in genes involved in vitamin and metabolite biosynthesis. These pseudogenization events nonetheless appear to be biased towards strains derived from soil-burrowing hosts, particularly the earliest-known underground transitions on the part of host lineages ca. 15 Mya (Beasley-Hall et al., 2021). Our findings also show genes encoding the alpha (*ureC*) and beta/gamma (*ureAB*) subunits of urease—responsible for nitrogen recycling, the key function of *Blattabacterium* for hosts—have been rendered non-functional or truncated in every soil-burrowing host lineage sampled, i.e., on at least six independent occasions.

We found evidence of positive selection in soil-burrowing host-derived strains in 12 of the 558 *Blattabacterium* genes examined here. However, none of these genes correspond to previously mentioned biosynthesis pathways that have been eroded through the pseudogenization or loss of other genes. Overall, these results indicate that soil-burrowing cockroach hosts are unable to produce most essential amino acids via their relationship with *Blattabacterium* and must acquire them from another source. There are two exceptions to this: cysteine is not thought to be provisioned by *Blattabacterium* as it is abundant in host hemolymph (Patiño-Navarrete et al., 2014), whereas proline is acquired by *Blattabacterium* from the host as the endosymbiont lacks a complete pathway for its synthesis (Sabree et al., 2009).

In addition to the loss of biosynthesis genes related to amino acids, we also observed the pseudogenization of genes encoding urease, involved in converting urea to ammonia. This enzyme is a critical component of the nitrogen recycling pathway in *Blattabacterium* and makes ammonia available to glutamate dehydrogenase (*gdhA*) to produce glutamate, a precursor to synthesis of other amino acids (Sabree et al., 2009). As we found no evidence for *gdhA* also being non-functional in any strains assessed here, it is possible the respective hosts can acquire ammonia from another source and/or have an alternate pathway for the conversion of urea in the first place. A similar phenomenon has been observed in the ant endosymbiont Blochmannia, in which urease is intact but the gene responsible for ammonia assimilation (glnA) is lost (Williams and Wernegreen, 2010). This disruption of a key step in the bacteria’s nitrogen recycling pathway means ammonia may be assimilated via alternate pathways in the endosymbiont, or by the host itself, to facilitate amino acid biosynthesis. Horizontal gene transfer is not known to occur from cockroach hosts to Blattabacterium, as in several other endosymbionts (Husnik et al., 2013; Kinjo et al., 2021; Sloan et al., 2014; Woolfit et al., 2009), but whether this phenomenon might occur in the reverse direction—for instance, in relation to nitrogen assimilation or amino acid biosynthesis—remains to be investigated. Alternate mechanisms facilitating nitrogen recycling in eroded *Blattabacterium* genomes deserve further analysis.

Differences in the extent of pseudogenization and loss of *Blattabacterium* genes within host clades are closely associated with the phylogenetic history of the Panesthiinae. Australian panesthiine cockroaches began transitioning underground approximately 15 million years ago (Mya), followed by multiple independent, and often broadly concurrent, colonisations of the subterranean environment from ca. 12 to 3 Mya (Beasley-Hall et al., 2021). The subset of soil-burrowing host lineages that do not display extensive *Blattabacterium* gene losses here are also those that made this transition relatively recently (Figure 1), implying that their endosymbiont genomes have not experienced analogous selection pressures for as long a period in evolutionary terms to facilitate genome erosions. Nonetheless, pseudogenizations are similarly more extensive in taxa from host lineages that diverged relatively early from wood-feeding lineages: *Macropanesthia kinkuna* str. BMK, present within lineage that diverged *ca*. 10 Mya from wood-feeders, displays a larger number of pseudogenization events than Macropanesthia *lithgowae* str. BML (*ca*. 5 Mya), for instance.

### 4.2 Trends in eroded biosynthesis pathways

Analogous biosynthetic pathways have been repeatedly and independently eroded in *Blattabacterium* symbionts of soil-burrowing cockroaches. The aphid co-obligate symbiont *Buchnera aphidicola* has also undergone the loss of genes associated with the synthesis of certain amino acids and vitamins in a stepwise manner, which may have been driven by the presence of the secondary symbiont *Serratia symbiotica* (Monnin et al., 2020) or vice versa (Manzano-Marín et al., 2016). Of particular interest is the loss in Buchnera of the ability to synthesise tryptophan, the only amino acid pathway that has been rendered non-functional in independent aphid lineages co-infected by *S. symbiotica*. This degradation has also occurred in the *Blattabacterium* strains presented here, and the pathway has been completely lost in genomes of CPU and MADAR as well as the *Carsonella* endosymbiont in psyllids (Sloan and Moran, 2012). Further trends emerge when considering the relationship between the energetic cost of pathways and their loss: tryptophan, phenylalanine, and methionine are some of the most expensive pathways in *Escherichia coli* in terms of the amount of ATP consumed (Kaleta et al., 2013), which might explain the complete loss of *metA* across *Blattabacterium* strains and the loss of *metB* and *pheA* in strains derived from soil-burrowing cockroaches. Leucine is also energetically expensive when measured by the proportion of glucose that its pathway demands from the glucose required for total amino acid production (Kaleta et al., 2013), and genes related to its synthesis have been lost in both the soil-burrowing members of the Panesthiinae and in strains CPU and MADAR (Fig 1). However, we note the cost of the production of these compounds may also be compensated for by their abundances in highly expressed proteins (Heizer et al., 2006).

Gene losses analogous to those observed here have also occurred in other sequenced *Blattabacterium* strains. Amino acid biosynthesis genes have been extensively lost in *Blattabacterium* from fast-evolving host lineages such as *Euphyllodromia* (Arab et al., 2020; Bourguignon et al., 2020; Kinjo et al., 2021), though these taxa have also experienced substantial gene loss in additional metabolic pathways such as coenzyme biosynthesis. Such losses appear to be associated with an elevated mutation rate in these strains as opposed to the parallel and selective losses seen in our ingroup (Bourguignon et al., 2020; Kinjo et al., 2021). The most similar losses to those seen in our dataset are in the termite *Mastotermes darwiniensis* str. MADAR and wood roach *Cryptocercus punctulatus* str. CPU (Kinjo et al., 2018). While these events have not occurred in the context of parallel evolution in their hosts, they have been putatively linked to concomitant changes to the host gut microbiome indirectly caused by host social behaviour, i.e., the direct feeding on hindgut fluids by offspring (proctodeal trophallaxis) (Tokuda et al., 2013). Our findings therefore raise the possibility that soil-burrowing lineages of the Panesthiinae may perform similar subsocial behaviours, as evidenced by their similarly eroded endosymbiont genomes, but this question is beyond the scope of the current study.

## 5 CONCLUSIONS

Here, we sequenced and examined 26 genomes of the obligate endosymbiont *Blattabacterium* from soil-burrowing and wood-feeding panesthiine cockroaches. Soil-burrowing species display parallel genome erosions and pseudogenization events that overwhelmingly affect genes involved in amino acid biosynthesis. We also observed pseudogenization of two genes encoding the subunits of urease, responsible for the first step in Blattabacterium’s key role as a nitrogen recycler for its cockroach hosts. These losses may be relatively predictable in that strongly-eroded pathways are associated with products that are very energetically costly to synthesise. Ultimately, our results indicate soil-burrowing cockroach hosts must acquire the building blocks of the relevant amino acids from alternate sources. *Blattabacterium* genome erosions nearly identical to those observed here have been documented in distantly related subsocial hosts, albeit without the pseudogenization of urease. As such, we speculate subsocial behaviours may facilitate the shrinkage of *Blattabacterium* genomes in soil-burrowing host lineages, e.g., the high-fidelity transfer of gut microbes with nutrient-provisioning functions via proctodeal trophallaxis.

Several outstanding questions have been raised from the results of this study. Future work on the system should ideally sample additional *Blattabacterium* genomes from soil-burrowing hosts, such as the relatives of *Geoscapheus dilatatus*—the species in which we observe the most extensive *Blattabacterium* erosions—to further assess varying degrees of genome shrinkage within and among host clades. The gut microbiome of the soil-burrowing cockroaches also remains to be characterised and evidence for a supplanting of endosymbiont functions by gut microbes may be uncovered if relevant functional shifts are observed in soil-burrowing vs. wood-feeding hosts. As the evolutionary trajectory of *Blattabacterium* in the soil-burrowing cockroaches appears to be somewhat predictable, our findings contribute to a wider narrative regarding the stochasticity of evolution as a whole. Gould (Gould, 1989) posited that if the “tape of life” were replayed, the organisms that arose might be unrecognisable because evolution is a random process in many aspects, including at the genetic level, and one might therefore expect organisms to have different genetic toolkits to overcome similar selective pressures. However, instances of parallel evolution have complicated this narrative. The present study contributes to the growing body of literature on genetic parallelism by demonstrating that the occurrence and extent of genome erosions in *Blattabacterium* are not only associated with parallel underground transitions in their hosts, but that the losses themselves have occurred in parallel and appear to be somewhat predictable or constrained.

## ACKNOWLEDGEMENTS

This research was supported by the Australian Research Council (grant no. FT160100463). The authors have no conflict of interest to disclose.

## DATA ACCESSIBILITY STATEMENT

All 25 annotated *Blattabacterium* genomes sequenced as part of this study will be uploaded to NCBI’s GenBank repository upon acceptance of this manuscript.

## AUTHOR CONTRIBUTIONS

PGBH and NL conceived the study. PGBH performed laboratory experiments, analysed the data, and wrote the manuscript. YK and CSPF helped with sequence data filtering and assembly. TB provided guidance regarding microbiome analysis. TGLK assisted with detecting gene losses. NL, HAR, JW, and SYWH helped to draft the manuscript. All authors edited the manuscript and approved the final draft.

