## Supplementary Material for "Shrinking in the dark: Parallel endosymbiont genome erosions are associated with repeated host transitions to an underground life"

### **1.1 | Host taxon sampling and *Blattabacterium* genome assemblies**

DNA extractions of fat body tissue from 100%-ethanol preserved cockroach specimens were performed using a QIAGEN DNeasy Blood and Tissue Kit. DNA sequencing was outsourced to Macrogen (Seoul, South Korea) or performed in-house at the Okinawa Institute of Science & Technology Graduate University using Illumina shotgun-sequencing technology, following the methods of (Kinjo et al., 2018).

Raw reads from our shotgun sequencing data were *de novo* assembled using SPAdes v.3.12.0 (Bankevich et al., 2012) with *k* values of 33, 55, 77, 91 and 121. SPAdes-derived contigs were then processed using TCSF-IMRA v.2.7.1 (Kinjo et al., 2015), a tool developed for the assembly of endosymbiont genomes. Briefly, TCSF was used to select contigs derived from *Blattabacterium* with homology searches against *Panesthia angustipennis yayeyamensis* str. BPAY (hereafter str. BPAY) (Kinjo et al., 2015), a close relative of the Australian Panesthiinae, using TBLASTX. We implemented this *a priori* hypothesis because gene rearrangements in *Blattabacterium* strains are extremely uncommon, with all known strains in the Panesthiinae showing high synteny (Kinjo et al., 2018; Neef et al., 2011; Patiño-Navarrete et al., 2013; Vicente et al., 2018). Host mitochondrial sequences were filtered out using previously published mitochondrial genomes for the group (Beasley-Hall et al., 2021). Second, IMRA was used to map raw reads onto TCSF-filtered contigs and iteratively reassemble and elongate them over 10 iterations with the same *k* values specified above. Scaffolds produced by IMRA were then processed using Chromosomer v.0.1.3 (Tamazian et al., 2016), which estimates the length between scaffolds and bridges them with Ns against a reference genome (str. BPAY). Finally, Pilon v.1.22 (Walker et al., 2014) was used to detect misassemblies using raw reads against our scaffolded assembly. We manually checked Ns inserted by Chromosomer that erroneously represented missing data using a whole-genome alignment computed with the progressiveMauve algorithm v.2015-02-25 (Darling et al., 2010) against str. BPAY.

To investigate the presence of potential secondary symbionts in these cockroaches, which may have also contributed to symbiont genome erosions in other insect hosts (Lamelas et al., 2011, 2008), raw reads for all non-pooled libraries were analysed using MetaPhlAn2 (Truong et al., 2015), which detects clade-specific marker genes from short read sequence data to perform taxonomic assignment and measures of abundance. Default settings and a 1% abundance detection threshold was selected for this analysis.

### **1.2 | Annotation and pseudogene detection**

Bacterial genome annotation was performed using Prokka v.1.14.5 (Seemann, 2014) with a similarity e-value cut-off of 1e-06, a minimum coverage of 80%, and a custom BLAST protein database constructed from all *Blattabacterium* genomes deposited via GenBank at the time of writing (Table S2). We used the Prokka implementations of ARAGORN (Laslett and Canback, 2004) to annotate tRNAs, Barrnap (<https://github.com/tseemann/barrnap>) for rRNAs, and Infernal+Rfam (Kolbe and Eddy, 2011) for ncRNAs. In addition to the 25 genomes generated here, we also annotated the previously published str. MACROPArhi from the host *Macropanesthia rhinoceros*, another soil-burrowing member of the Panesthiinae (Bourguignon et al., 2020).

Gene loss was assessed using Prokka annotations and manually validated using a progressiveMauve alignment against str. BPAY; genes were considered lost if the relevant sequence was completely missing relative to BPAY and was flanked on either side by a contiguous sequence—that is, the annotation was not missing or disrupted due to missing data bridging scaffolds. BLASTN was used to double-check the absence of genes by querying sequences from closely related strains against the genome in question using the Megablast algorithm with a word size of 28 and e-value 1e-10. Orthologous genes were inferred from our Prokka-annotated genomes using OMA v2.5.0 (Altenhoff et al., 2021) with default settings, which were then processed using translatorX (Abascal et al., 2010) to yield 715 orthologous groups (hereafter OGs) aligned by codon position. Pseudogene detection was performed on OGs using Pseudofinder, a program designed for the inference of pseudogenization events in prokaryotic genomes, using default settings (Syberg-Olsen et al., 2022). We used the same *Blattabacterium* database as in our Prokka annotation step and an 80% truncation cut-off such that a gene over 20% shorter than the average length of its BLAST hits would be considered a putative pseudogene. We used this candidate set for the selection analyses detailed below. Finally, given the fact that certain endosymbionts can correct frameshift mutations—meaning apparent pseudogenes may have functional ORFs (Tamas et al., 2008)—and the fact that the majority of genomes presented here were not closed, we manually validated pseudogenised genes relevant to the biosynthesis of amino acids, vitamins, and metabolites (Figure 1) against the *Blattabacterium* genome of the relevant taxon using the Megablast algorithm of BLASTN with a word size of 28 and e-value 1e-10.

### **1.3 | Tests of selection**

OGs were filtered individually such that each host species was not represented more than once in the alignment and intraspecific symbiont sequences were excluded (e.g., multiple *Blattabacterium* strains from *Geoscapheus dilatatus*). Putative pseudogenes identified by Pseudofinder were individually removed from each OG prior to analysis. Alignments were also excluded from the analysis if they did not correspond to 1:1 orthologs of protein-coding genes, only contained *Blattabacterium* strains from a single host group (e.g., exclusively soil-burrowing hosts), or if the respective OG contained less than 10 taxa, representing less than 50% of the above number of hosts after intraspecific data was excluded. Our final dataset for downstream analysis consisted of 578 OGs. The UniProt database (UniProt Consortium, 2021) was used to validate Prokka-annotated gene abbreviations and protein names for consistency and retrieve Gene Orthology terms for each OG. GWideCodeML (Macías et al., 2020) was used to prepare trimmed phylogenies and control files for each OG prior to selection analysis.

Tests for signatures of relaxed and positive selection were conducted using the *codeml* program in PAML v.4 (Yang, 2007). *Blattabacterium* strains originating from soil-burrowing hosts were selected as lineages of interest (foreground branches) for the analysis. *codeml* makes use of nonsynonymous/synonymous substitution ratios (dN/dS) to measure the mode and strength of selection acting on genes. Depending on the model selected, *codeml* permits dN/dS ratios to vary at different sites within the same gene (site models), at different branches in the phylogeny (branch models), or at particular sites on particular branches (branch-site models). An increase in dN/dS can indicate either positive selection or a relaxation of purifying selection, though the majority of non-synonymous changes across an entire gene are assumed to be either selectively slightly deleterious or neutral (Hughes, 2007; Kimura, 1968; Ohta, 1973). As such, we operated under the assumption that a significant increase in dN/dS over an entire gene is indicative of relaxed selection, whereas an elevated dN/dS for a certain number of sites within a gene, where beneficial mutations have occurred, is indicative of positive selection. With this in mind, we used branch models to assess relaxed selection and branch-site models to assess positive selection (Mitterboeck et al., 2017). In the first case, we selected the one ratio (M0) and two ratio (M1) branch models, which enforce a single dN/dS for the entire phylogeny or allow the ratio to differ between foreground and background branches, respectively. To detect signatures of positive selection we used the branch-site models A1 and A, the latter of which allows sites to have dN/dS values of >1. Likelihood ratio tests were conducted separately for each gene to assess significant dN/dS differences between foreground and background branches. Resulting *p* values were corrected using the Benjamini-Hochberg procedure with a false discovery rate of 0.05 (Benjamini and Hochberg, 1995).

### **2 | Supplementary tables**

**TABLE S1**  Soil-burrowing and wood-feeding panesthiine cockroach hosts used in this study. All taxa are endemic to the Australian mainland and are soil-burrowers except for *Panesthia*. QLD = Queensland; VIC = Victoria; NSW = New South Wales; SA = South Australia; WA = Western Australia.

| **Genus** | **Species** | **Collection locality** | **GenBank accession** |
| --- | --- | --- | --- |
| *Geoscapheus* | *crenulatus crenulatus* | Rainbow Beach, QLD | TBA |
|  | *dilatatus* | Chinkapook, VIC | TBA |
|  |  | Gilgandra, NSW | TBA |
|  |  | Mitchell, QLD | TBA |
|  |  | Patchewollock, VIC | TBA |
|  |  | Renmark, SA | TBA |
|  |  | Walpeup, VIC | TBA |
|  |  | Yenda, NSW | TBA |
|  | *robustus* | Laverton, WA | TBA |
|  | *woodwardi* | Muttaburra, QLD | TBA |
| *Macropanesthia* | *kinkuna* | Coonarr, QLD | TBA |
|  | *lithgowae* | Nudley State Forest, QLD | TBA |
|  | *mutica* | Dinden State Forest, QLD | TBA |
|  | *rothi* | Agnes Water, QLD | TBA |
|  | *rhinoceros* | - | CP059200.1 |
| *Neogeoscapheus* | *dahmsi* | Taroom, QLD | TBA |
|  | *hanni* | Mt. Molloy, QLD | TBA |
| *Parapanesthia* | *gigantea* | Warwick, QLD | TBA |
| *Panesthia* | *australis* | Wiseleigh, VIC | TBA |
|  | *obtusa* | Mt. Blackwood, QLD | TBA |
|  | *sloanei* | Mossman Gorge, QLD | TBA |
|  | *tryoni tegminifera* | Mt. Lewis, QLD | TBA |
|  |  | Dorrigo National Park, QLD | TBA |
|  | *tryoni tryoni* | Kroombit Tops National Park, QLD | TBA |
|  |  | Eungella, QLD | TBA |
|  |  | Sarina, QLD | TBA |

**TABLE S2**  Database of published *Blattabacterium* strains supplied to Prokka for annotation of genomes sequenced here.

| ***Blattabacterium* strain name** | **Host species** | **GenBank accession** |
| --- | --- | --- |
| ALLACaus | *Allacta australiensis* | CP059223 |
| ALLACbim | *Allacta bimaculata* | CP059221 |
| AMAZOsp | *Amazonina* sp. | CP059220 |
| ANALLAmet | *Anallacta methanoides* | CP059219 |
| ANAPcal | *Anaplecta calosoma* | CP059218 |
| ANAPome | *Anaplecta omei* | CP059217 |
| BALTAsp | *Balta* sp. | CP059216 |
| BEYBkur | *Beybienkoa kurandanensis* | CP059215 |
| BGE | *Blattella germanica* | CP001487 |
| BGIGA | *Blaberus giganteus* | CP003535 |
| BNCIN | *Nauphoeta cinerea* | CP005488 |
| BPAA | *Panesthia angustipennis spadica* | AP012548 |
| BPAY | *Panesthia angustipennis yayeyamensis* | AP014609 |
| BPLAN | *Periplaneta americana* | CP001429 |
| CARBpar | *Carbrunneria paramaxi* | CP059213 |
| CCLhc | *Cryptocercus punctulatus* | CP029844 |
| CHORISOsp | *Chorisoserrata* sp. | CP059214 |
| CKYod | *Cryptocercus punctulatus* | CP029820 |
| COSMOsp | *Cosmozosteria* sp. | CP059212 |
| CPU | *Cryptocercus punctulatus* | CP003015 |
| CPUbr | *Cryptocercus punctulatus* | CP029816 |
| CPUbt | *Cryptocercus punctulatus* | CP029813 |
| CPUmc | *Cryptocercus punctulatus* | CP029815 |
| CPUml | *Cryptocercus punctulatus* | AP014610 |
| CPUmp | *Cryptocercus punctulatus* | CP029814 |
| CPUpc | *Cryptocercus punctulatus* | CP029811 |
| CPUsm | *Cryptocercus punctulatus* | CP029810 |
| CPUsv | *Cryptocercus punctulatus* | CP029812 |
| CPUwf | *Cryptocercus punctulatus* | CP029818 |
| CYRTOsp | *Cyrtotria* sp. | CP059211 |
| DEROpau | *Deropeltis paulinoi* | CP059210 |
| DPU | *Diploptera punctata* | CP049785 |
| DUCHAsp | *Duchailluia* sp. | CP060246 |
| DYAKIkur | *Dyakinodes kurandensis* | CP059209 |
| ECTOBIsp | *Ectobius* sp. | CP059208 |
| ECTONUhan | *Ectoneura hanitschi* | CP059207 |
| EPILAmay | *Epilampra maya* | CP059206 |
| ESCALves | *Escala vestjensi* | CP059205 |
| EUPHYsp | *Euphyllodromia* sp. | CP059204 |
| EUPOLsin | *Eupolyphaga sinensis* | CP059203 |
| GROMPgra | *Gromphadorhina grandidieri* | CP059202 |
| GYNAcap | *Gyna capucina* | CP059201 |
| LAMPROsp | *Lamproblatta* sp. | CP060245 |
| MACROPArhi | *Macropanesthia rhinoceros* | CP059200 |
| MADAR | *Mastotermes darwiniensis* | CP003000 |
| MEDIASdel | *Mediastinia* sp. | CP059199 |
| MEGALOsp | *Megaloblatta* sp. | CP059198 |
| MELANOZsp | *Melanozosteria* sp. | CP059197 |
| METHAsp | *Methana* sp. | CP059196 |
| NEOLAXmac | *Neolaxta mackerrasae* | CP059195 |
| NYCTIBsp | *Nyctibora* sp. | CP059225 |
| OPISTHori | *Opisthoplatia orientalis* | CP059194 |
| PANESsp | *Panesthia* sp. | CP059193 |
| PARANAUcir | *Paranauphoeta circumdata* | CP059192 |
| PARATEMsp | *Paratemnopteryx* sp. | CP059191 |
| PARCOBvir | *Parcoblatta virginica* | CP059190 |
| PHYLLODsp | *Phyllodromica* sp. | CP059189 |
| PLATYZsp | *Platyzosteria* sp. | CP059188 |
| POLYPHAGsp | *Polyphagoides* sp. | CP059187 |
| PROTAGlug | *Protagonista lugubris* | CP059224 |
| RHABDOBsp | *Rhabdoblatta* sp. | CP059186 |
| SCHULTlam | *Schultesia lampyridiformis* | CP059185 |
| SHELFORDElat | *Shelfordella lateralis* | CP059184 |
| SHELFORDIsp | *Shelfordina* sp. | CP059183 |
| STAT | *Salganea taiwanensis taiwanensis* | AP014608 |
| Tarazona | *Blatta orientalis* | CP003605 |
| THEREAreg | *Therea regularis* | CP059182 |
| TRYONIpar | *Tryonicus parvus* | CP059181 |

**Supplementary File S1:** Database of orthologous groups from OMA, including Gene Ontology terms.

**Supplementary File S2:** Pseudofinder output, showing all inferred instances of putative pseudogenization in examined *Blattabacterium* genomes. These candidates were refined downstream via manual curation.

**Supplementary File S3:** Results of genome-wide relaxed selection analyses in PAML.

**Supplementary File S4:** Results of genome-wide positive selection analyses in PAML.

**Supplementary File S5:** *ureAB* alignment.

**Supplementary File S6:** *ureC* alignment.
